# Functional coupling between the α4 – α5 loop and allosteric site on MKP5 is a critical determinant of catalysis

**DOI:** 10.64898/2026.06.01.729370

**Authors:** Ramu Manjula, Lubna Ghanem, Erin Skeens, Leanna Bai, George P. Lisi, Elias J. Lolis, Anton M. Bennett

**Affiliations:** Department of Pharmacology, Yale University School of Medicine, New Haven, Connecticut, USA; Department of Molecular Biology, Cell Biology and Biochemistry, Brown University, Providence, Rhode Island, USA; Yale Center for Molecular and Systems Metabolism, Yale University School of Medicine, New Haven, Connecticut, USA

**Keywords:** Protein tyrosine phosphatase, mitogen-activated protein kinase, allosteric site

## Abstract

Dual-specificity phosphatases (DUSPs) that inactivate mitogen-activated protein kinases (MAPKs) are called MAPK phosphatases (MKPs). The stress-responsive MKP, MKP5, dephosphorylates p38 MAPK and c-Jun NH_2_-terminal kinase (JNK). Previously, we identified an allosteric compound that binds to the site within the phosphatase domain of MKP5 that is critical for the binding and dephosphorylation of both p38 MAPK and JNK. The allosteric site, comprised of the α4-α5 loop, is an essential region for transmitting MAPK binding to the catalytic site. Here, we examine the contribution of additional structural rearrangements that occur upon allosteric site engagement. We show that binding of an inhibitor that engages the MKP5 allosteric site induces conformational changes in the α4-α5 loop. Mutants of residues in this loop inhibited enzymatic activity, and some mutants exhibited changes in dynamics, indicating that this loop has a significant structural and dynamic role in controlling MKP5 catalysis. Enzymatic and NMR studies supported the interpretation that conformational changes and dynamics in the α4-α5 loop are required for enzymatic function. Additionally, alanine mutants of R442 (α4-α5 loop), the catalytic base D377 (β4-α2 loop), and Q409A and S413A (β5-α3 loop) disrupted catalytic activity. These results highlight the α4-α5, β4-α2, and β5-α3 loops as structurally, dynamically, and functionally interconnected for communication between the allosteric and enzymatic sites.

## Introduction

Protein phosphorylation is a reversible process that is regulated by protein kinases and protein phosphatases (1). The dual-specificity family of protein phosphatases (DUSPs) constitutes a sub-family within the family of protein tyrosine phosphatases (2,3). A sub-set of DUSPs, known as the mitogen-activated protein kinase (MAPK) phosphatases (MKPs) exhibit unique substrate selectivity to the MAPKs (3,4). There are 10 active MKPs, each endowed with the capacity to specifically dephosphorylate the MAPKs on phospho-threonine/serine and phospho-tyrosine residues within the activation loop of the MAPKs, resulting in their inactivation (5–7). The MKPs mediate MAPK dephosphorylation through the engagement of an essential cysteine residue located within the catalytic domain and subsequent generation of a thiophosphoryl intermediate (7). A conserved aspartate residue located within loop β4-α2 of the enzymatic site is essential for the transfer of the phosphotyrosyl group from the substrate to the enzyme through protonation of the leaving group (8). The reaction is terminated upon hydrolysis of the phosphoenzyme, which is catalyzed by the essential aspartate residue that serves as a general base (7,9).

Although much has been learned about the molecular basis of MKP-mediated dephosphorylation through MAPK activation loop catalysis, a complete understanding of the regulatory determinants of the MKPs still remains unresolved. The importance of providing an in-depth molecular understanding of the mechanisms of MAPK-mediated dephosphorylation is highlighted by the complexity in which MKPs elicit downstream signaling. The MAPKs regulate many vital cellular processes through serine/threonine phosphorylation, and their dysregulation can lead to cancer, immunological, and metabolic diseases (10,11). The MAPKs elicit both positive and negative signaling effects, and their inactivation by the MKPs results in complex physiological and pathophysiological outcomes that are an integration of their net activities (4,12–14).

We identified an allosteric site within the MKP5 catalytic domain (CD) that provided further insight into the regulation of the MKPs. The allosteric site represents an interface that coordinates the reorganization of critical residues around the catalytic cysteine that are required for catalysis (15,16). The structural integrity of the MKP5 allosteric site is maintained by several hydrophobic residues that also influence the entire phosphatase domain. Engagement of a small-molecule MKP5 inhibitor at the allosteric site is responsible for extensive loop movements but does not affect the conformations of C408 or the catalytic base D377 (16). Specifically, the shift in the α4-α5 loop of MKP5 impacts the orientation of the β5-α3 loop, which forms the catalytic site. Insight from studying the structural changes that occur upon engagement of the allosteric site revealed that residues ^445^ISPN^448^ in the α4-α5 loop shift the peptide backbone. In this scenario, we hypothesized that the allosteric loop (α4-α5 and Fig. S1) and the catalytic loops (β5-α3, β4-α2 and Fig. S1) are important for mediating MKP5 catalysis. Given the importance of the MKP5 allosteric site to the regulation of phosphatase activity, MAPK binding and the conservation of this site in other active MKPs, we reasoned that further investigation into the molecular interactions that reside around the allosteric site will enhance our understanding of MKP5 and, therefore, MKP regulation more broadly.

Here, we demonstrate that maintaining the integrity of the α4-α5 loop, which contributes to the allosteric pocket, is essential for its connection to the catalytic loops (β5-α3 and β4-α2). X-ray crystallography and NMR support the interpretation that, in addition to the requirement for allosteric hydrophobicity, the α4-α5 loop conforms to accommodate allosteric site occupancy to promote MKP5 catalysis. Taken together, these studies extend the structural landscape within the MKP5 catalytic domain that regulates its activity.

## Results

### Essential role of α4-α5 loop for MKP5 catalytic activity

Previously, we reported that Compound 1 (Cmpd 1; 3,3-dimethyl-1-((9-(methylthio)-5,6-dihydrothieno[3,4-h]quinazolin-2-yl)thio)butan-2-one) binding to the allosteric site of MKP5, induces a conformational shift of the α4-α5 loop (Fig. S1 *A, B,* S2) (16). The α4-α5 loop is comprised of residues ^445^ISPN^448^. To test the contribution of the α4-α5 loop residues to regulate MKP5 catalytic activity, we generated I445A, S446G, P447V and N448A variants within this loop and measured their enzymatic activity using substrates p-nitrophenyl phosphate (pNPP), p38 MAPK phosphopeptide and 6,8-difluoro-4-methylumbelliferyl phosphate (DiFMUP) (Fig. 1). The variant I445A was found to be significantly reduced by ∼50% in activity as compared with WT (Fig. 1*A*). Additionally, the activity of P447V and N448A were also significantly inhibited by ∼90% and ∼80%, respectively as compared with WT (Fig. 1*A*). In contrast, S446G was only impaired in its activity by ∼20% as compared with WT (Fig. 1*A*). We observed similar effects of these variants using DiFMUP as a substrate with P447V similarly demonstrating the most deleterious reduction in activity (Fig. 1*B*). Although the variants I445A and N448A using DiFMUP were of the identical rank order as compared with the *p*NPP substrate, at saturating DiFMUP concentrations, both exhibited comparable activities (Fig. 1*B*). Next, we measured catalytic activity of these variants against the p38 MAPK phosphopeptide (Fig. 1*C*). A similar rank order of inhibition was identified with S446G retaining much of its activity followed by I445A>P447V=N448A as compared with WT (Fig. 1*C*).

**Figure 1.**
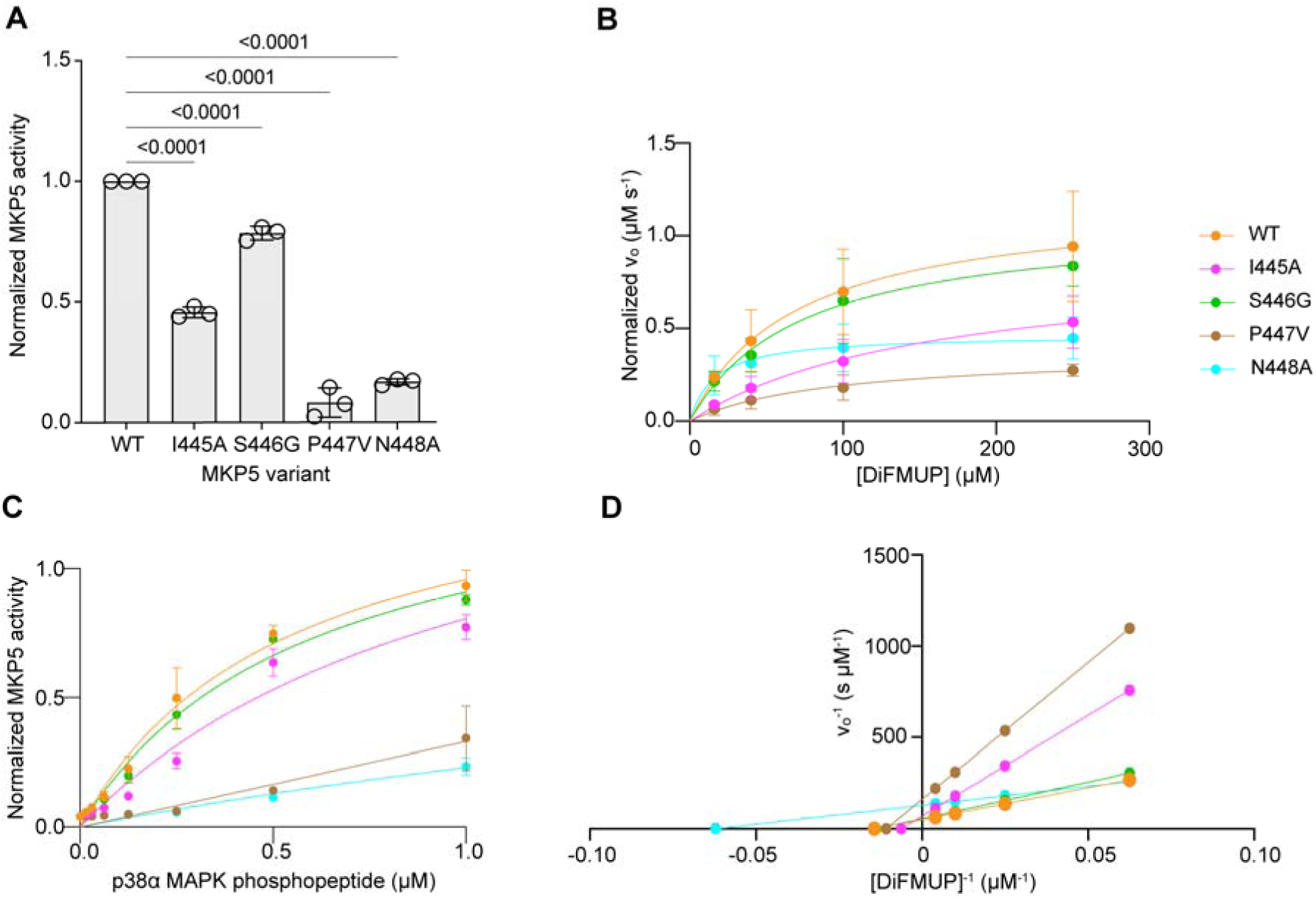
Effect of MKP5 α4-α5 loop on phosphatase activity. MKP5 WT and α4-α5 loop mutant proteins were assayed for phosphatase activity using *A*, *p*NPP (1.25 mM) for 10 min at 37°C. Absorbance was measured at 405 nm. Significance was determined by one-way ANOVA. *B*, MKP5 WT or α4-α5 loop mutant proteins were incubated with a concentration gradient of DiFMUP. Data were fit to the Michaelis-Menten equation. *C,* 1μM MKP5 WT and α4-α5 loop mutant proteins were incubated with p38 MAPK phosphopeptide for 20 min at 30°C. Malachite green was added and incubated at room temperature for 15 min. Absorbance was measured at 620 nm. Data were fit to the Michaelis-Menten equation. *D*, Lineweaver-Burk analysis of the DiFMUP kinetic assay performed in *B*.

To further define the kinetic parameters of these MKP5 variants, we measured their initial velocities using DiFMUP as a substrate and derived *V_m_*_ax_, *K_m,_* and *k*_cat_ values by direct curve-fitting of the data to the Michaelis-Menten equation (Fig. 1*D* and Table 1). Consistent with our previously reported observations, MKP5 WT exhibited a *K*_m_ = 69.9 μM, *k*_cat_ = 0.2 s^-1^ and *V*_max_ = 0.02 μM.s^-1^ (Fig. 1*D* and Table 1). The MKP5 variant S446G showed similar kinetic parameters as compared with WT, consistent with its modest reduction in steady-state activity (Fig. 1*D* and Table 1). However, I445A and P447V showed impaired *V_m_*_ax_ values of 0.015 μM.s^-1^ and 0.006 μM.s^-1^, respectively, as well as increased *K_m_* and reduced *k*_cat_ values (Fig. 1*D* and Table 1). Interestingly, N448A showed reduced *V*_max_ (0.008 μM.s^-1^) and *k*_cat_ (0.08 s^-1^) despite exhibiting a *K_m_* that was markedly decreased (16.1 μM) as compared with WT (Fig. 1*D* and Table 1). We calculated the substrate specificity constant, *k*_cat_/*K_m_* (μM^-1^ s^-1^) to assess whether these variants impacted selectivity towards DiFMUP (Table 1). S446G showed little difference in *k*_cat_/*K_m_*, whereas I445A and P447V variants had ∼2-3-fold lower *k*_cat_/*K_m_* for DiFMUP as compared with WT (Table 1). The *k*_cat_/*K_m_* of N448A (4.8 x 10^-3^ μM^-1^ s^-1^) was greater than that of WT (2.86 x 10^-3^ μM^-1^ s^-1^) (Table 1). Taken together, these data indicate that residues ^445^ISPN^448^ within the α4-α5 loop contribute directly to the catalytic activity and/or substrate affinity of MKP5.

**Table 1:** Kinetic parameters of MKP5 α4–α5 loop variants.

|  | **WT** | **I445A** | **S446G** | **P447V** | **N448A** |
| --- | --- | --- | --- | --- | --- |
| $k_{cat}$ (s<sup>-1</sup>) | 0.200 | 0.145 | 0.181 | 0.062 | 0.078 |
| $K_m$ (μM) | 69.84 | 160.20 | 72.71 | 92.95 | 16.08 |
| $V_{max}$ (μM.s<sup>-1</sup>) | 0.020 | 0.015 | 0.018 | 0.006 | 0.008 |
| $k_{cat}/K_m$ (μM<sup>-1</sup>.s<sup>-1</sup>) | 2.86 x 10<sup>-3</sup> | 9.07 x 10<sup>-4</sup> | 2.49 x 10<sup>-3</sup> | 6.67 x 10<sup>-4</sup> | 4.82 x 10<sup>-3</sup> |

### NMR reveals crosstalk between the MKP5 **α**4-**α**5 loop and catalytic site

To further assess the structural basis for the effects of the variants on MKP5 activity we used NMR. We generated variants I445A, S446G, P447V, and N448A and noted that each induced significant chemical shift perturbations surrounding the site of mutation within the loop (Fig. 2). Despite these local changes to the MKP5 CD structure, its overall fold (based on resonance dispersion in ^1^H-^15^N HSQC spectra) and structural stability (via thermal unfolding monitored by CD spectroscopy) was unaffected (Figs. S3-S7). Additionally, P447V and N448A MKP5-CD also exhibit significant chemical shift perturbations at the catalytic residue D377 and at other residues on the D377-containing loop, suggesting a high degree of crosstalk and sensitivity to these mutations at the catalytic site (Fig. 2).

**Figure 2.**
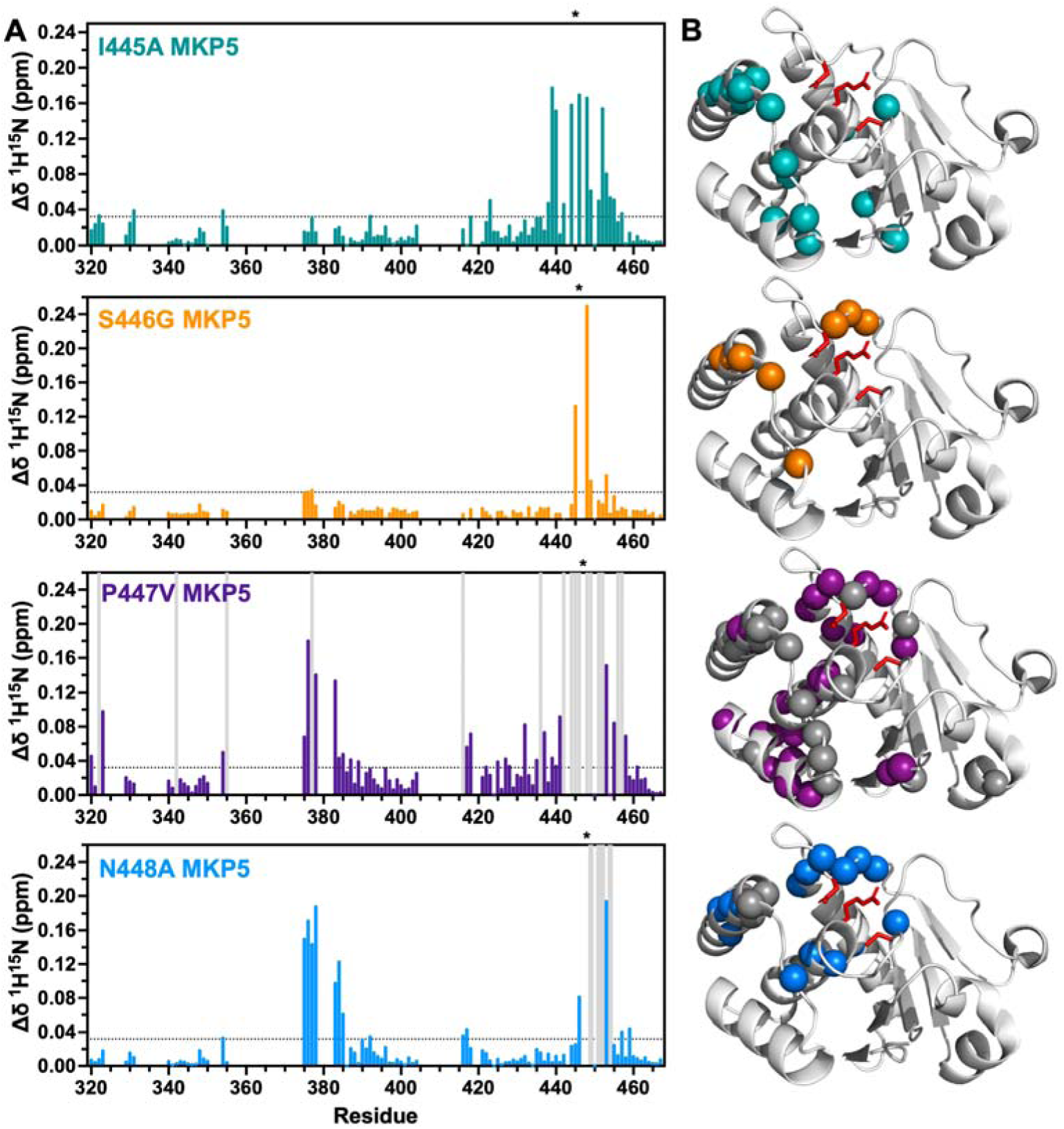
Structural impact of MKP5 loop variants. *A*, Chemical shift perturbation plots for I445A (cyan), S446G (orange), P447V (purple), and N448A (blue) MKP5-CD. Colored bars represent ^1^H^15^N combined chemical shift perturbations (Δδ), calculated for each variant relative to WT MKP5-CD. Chemical shift perturbations significance was defined as1.5σ above the 10% trimmed mean of all data sets (dashed horizontal line). Grey vertical lines denote resonances that exhibit line broadening and the site of mutation is noted by an asterisk above each plot. *B,* Significant chemical shift perturbations are plotted onto the MKP5-CD structure (PDB: 2OUD) in cyan (I445A), orange (S446G), purple (P447V), and blue (N448A) spheres. Line-broadened resonances are indicated by gray spheres and the side chains of catalytic site residues (D377, C408, R414) are highlighted in red.

The NMR data correlate well to kinetic experiments, where P447V and N448A have the lowest catalytic activities relative to WT MKP5 (Fig. 1). These variants are the only pair to induce exchange broadening in the NMR spectra, which is most prominent in P447V following removal of the bulky proline residue. Line broadening surrounds the cluster of large NMR chemical shift perturbations and suggests a change in flexibility near the catalytic site and allosteric pocket. Critically, the structural effect of the P447V and N448A mutations, based on strong NMR chemical shift perturbations, propagates to the catalytic site, while I445A and S446G variants, which retain catalytic functions similar to WT MKP5, show weak NMR chemical shift perturbations beyond the site of mutation (Fig. 2). These data point to critical positions within the MKP5 allosteric loop that affect its atomic structure and dynamics to engage in molecular crosstalk with the active site.

### Structural comparison of apo-N448A and Cmpd 1-N448A complexes for MKP5 catalysis

The WT MKP5-CD was previously demonstrated to undergo conformational changes in two loops upon binding of the allosteric inhibitor Cmpd 1 (Fig. S2). One loop consists of residues ^445^ISPN^448^ (α4-α5 loop), and the other comprises residues ^410^AGVS^413^ (β5-α3 loop), which causes MKP5 to lose catalytic activity when disrupted. To further investigate the structural impact of residues ^445^ISPN^448^, efforts were made to crystallize the four apo- and Cmpd 1-bound mutants and compare them to the apo-WT and Cmpd 1-WT structures to elucidate the mechanistic role of each residue in the ^445^ISPN^448^ loop (Table S1). Although we crystallized the four mutants, only N448A was determined in both the apo and allosteric Cmpd 1-bound forms, providing comprehensive insight into its role in enzymatic activity and inhibition. The other mutants were crystallized only in apo or Cmpd 1-bound structures.

The N448A mutation significantly reduces catalytic activity, exhibiting the lowest among the four mutants against the p38α phosphopeptide substrate (Fig. 1*C*). To examine the structural basis for decreased enzymatic activity of apo-N448A, its structure was compared to the active apo-WT MKP5-CD (Fig. 3*A*). A significant change was observed at the mutation site within the α4-α5 loop, confirming the NMR dynamics of apo-N448A, along with notable structural alterations at the β4-α2 loop where D377 is situated (Fig. 3*A*). Dynamics also occurs by the mutation breaking a network of strong hydrogen bonds among the side chain atoms of N448, D377, and S413. These atoms are located within loops α4-α5 and β4-α2, as well as at the end of the α3 helix next to β5-α3 (Fig. 3*B*). In addition to residue-level changes, the disruption prompts a 4 Å movement of the entire β4-α2 loop, which rotates the D377 side chain almost 180°. This shifts the initial position of D377 relative to the α4-α5 and β5-α3 loops within WT MKP5-CD to an orientation where D377 faces away from these loops in the N448A mutation. The movement and rotation of D377 cause its position to shift by 9 Å from the original WT structure. The distance of D377 to S413 and to the mutated N448A increased to 11 Å and 12 Å, respectively. These substantial changes in apo N448A MKP5-CD highlight the essential role of the wild-type residues in the α4-α5, β5-α3, and β4-α2 loops in maintaining structural stability and catalytic activity through side chain hydrogen bonding (Fig. 3*C*) and are consistent with the NMR dynamics of this mutation (Fig. 2).

**Figure 3.**
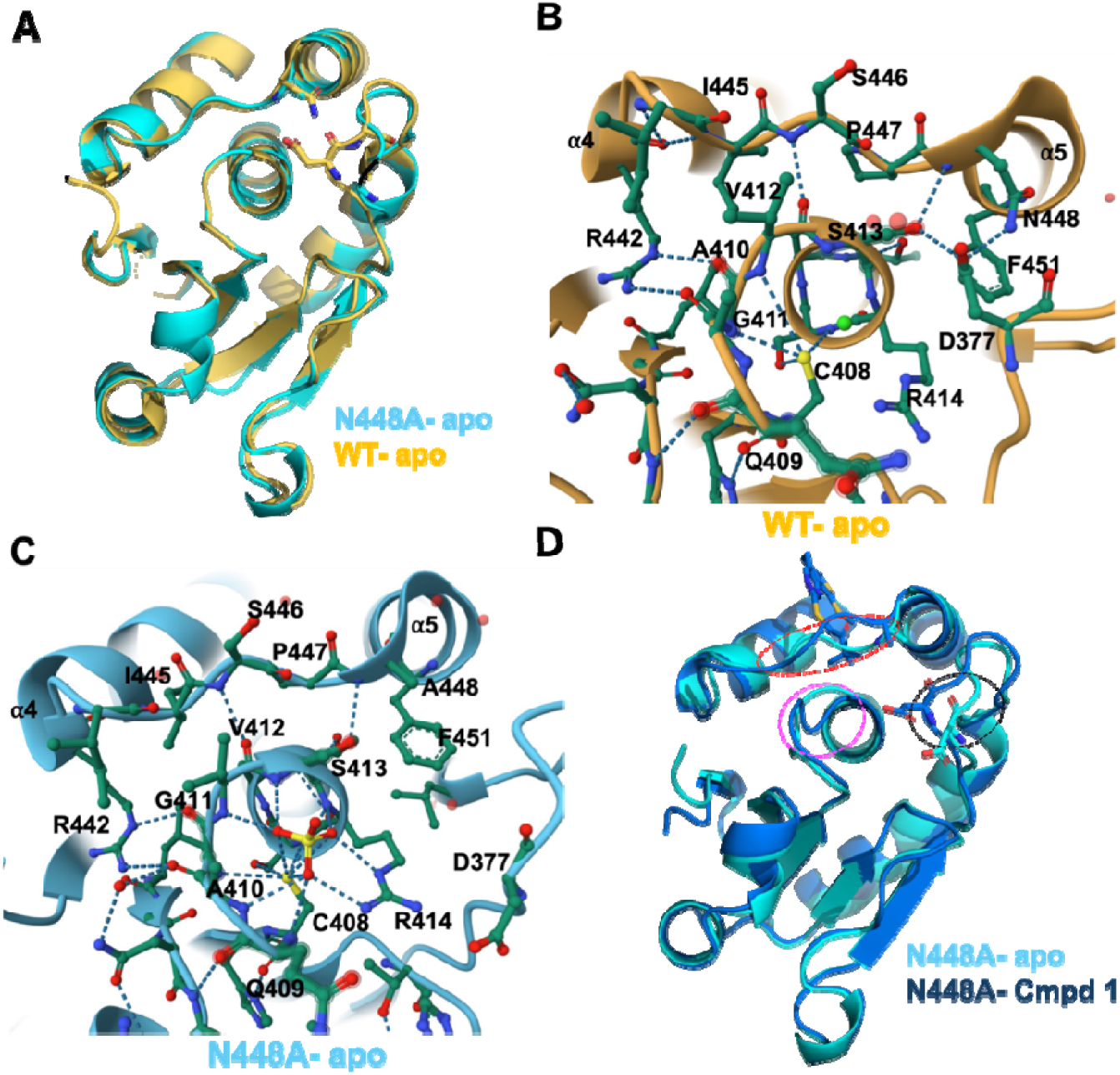
Structural comparison of WT and N448A MKP5-CD structures. *A*, Superposition of apo-WT (wheat) and apo-N448A (cyan) structures, showing high overall similarity. In apo-WT MKP5, the side chains of N448 and D377 form a 2.7 Å hydrogen bond (orange dots). In the apo-N448A structure, the distance to D377 is 12 Å (cyan arrows), with a conformational change in the β4-α2 loop and D377. *B*, Close-up view of the active-site hydrogen-bonding network of apo-WT MKP5-CD (2OUD). *C*, Corresponding view of the inactive apo-N448A mutant highlighting the disrupted hydrogen-bonding network involving N448A, D377, and S413 that is essential for enzymatic activity in the WT MKP5-CD. *D*, Superposition of apo N448A (cyan) and Cmpd1-N448A (blue) structures. Apo-N448A has an extra loop (β4-α2) due to a different orientation of D377 compared to Cmpd 1-bound N448A. The loops are the α4–α5 loop (red) next to Cmpd 1, the β5-α3 loop (magenta), and the β4-α2 loop (black), which form the catalytic pocket. Hydrogen bonds are shown as dashed lines, and key residues are displayed in stick representation. The figures were generated using the PyMol (*A* & *D*) and Mol* (*B* & *C*) molecular visualization tools.

We examined the three-dimensional structures of Cmpd 1-WT MKP5 CD and Cmpd 1-bound N448A to elucidate the mechanisms of inactivation in greater detail. At high concentrations during crystallization, the asymmetric unit contains six independent Cmpd 1-N448A complexes, the same as the Cmpd 1-WT MKP5 CD when it was first crystallized (16). The RMSD of Cα atoms across the Cmpd 1 structures ranges from 0.16 to 0.30 Å, indicating no significant difference. In contrast to the Cmpd 1-WT MKP5 CD, a local variation is observed at each of D377 among the six side chains of Cmpd 1-N448A. The D377 side chains adopt different orientations, with rotations and distances ranging from 0° to 135° relative to the superimposed structures of D377 of Cmpd 1-WT MKP5-CD (Fig. S8A). Only one orientation of the mutant’s D377 aligned structurally with the Cmpd 1-WT MKP5-CD (at 0° rotation), but surprisingly, it lacked the hydrogen bond present solely in the side chains of D377 and N448 in the WT MKP5-CD (Fig. S8B). As the rotation of the mutant D377 side chain increases, the distance from the compared Cmpd 1-WT MKP5-CD expands from 1.5 Å to 6.8 Å in different conformations (Fig. S8C-G). A maximum rotation of 135° in the crystal is paired with an extra 2 Å conformational change of the entire β4-α2 loop (Fig. S8A, S8G). All of these N448A mutant orientations are due to the weak van der Waals interactions that substitute for the WT side-chain hydrogen bonds between D377 and N448. It is the weak van der Waals interactions that allow different D377 conformations in each of the six mutant proteins in the crystal. We also measured the distance between the D377 side-chain carboxylate and the N448A methyl group for each mutant protein. The WT hydrogen-bond distance between the side chain atoms is 2.7 Å (Fig. S9A), while the six D377-N448A mutants range from 4.7 Å to over 11 Å, with an average of 8.2 Å (Fig. S9B-G). These distances to N448A are determined by weak van der Waals interactions of D377 and the amino acids H357, Q383, S413, R414, and F451 at various positions, reflecting the dynamic behavior of D377 in the N448A mutation.

We also analyzed the structures of Cmpd 1-N448A and apo-N448A to determine any structural effects of binding Cmpd 1 or the N448A mutation (Fig. 3*D*). In apo-N448A MKP5-CD, the disappearance of the hydrogen bond between N448 and D377 causes the β4-α2 loop to shift by 3 Å. This movement leads to further conformational changes in the α4-α5 and β5-α3 loops, resulting in a rearrangement of all three loops that compromises the active-site geometry (Fig. 3*D*). The Cmpd 1-N448A complex exhibits a more limited pattern of structural perturbation compared to apo-N448A. While conformational changes are still observed in the α4-α5 and β5-α3 loops, the β4-α2 loop in Cmpd 1-N448A surprisingly adopts a conformation comparable to that of both apo- and Cmpd 1-bound WT MKP5-CD. This indicates that the effects of Cmpd 1 binding partially offset the destabilization caused by the N448A mutation driven by van der Waals forces. Taken together, these findings highlight the critical role of N448 in the α4-α5 loop, and of the catalytic loops β4-α2 and β5-α3, in maintaining the integrity of the MKP5 PTP domain. The hydrogen-bonding network involving D377, S413, and N448 is crucial for maintaining the structural stability of the catalytic pocket. Disrupting this network—either through the mutation N448 alone or combined with Cmpd 1 binding causes some residues in the ^445^ISPN^448^, ^410^AGVS^413^, and ^376^TDSN^379^ loops to shift more than 10 Å from their positions in the wild type protein.

### Conformational changes in **β**4-**α**2, **β**5-**α**3 and **α**4-**α**5 induced by allosteric site engagement regulate MKP5 catalysis

We observed that of the remaining three variants, I445A was crystallized only in its apo form, whereas S446G and P447V mutants were crystallized in complex with Cmpd 1 (Table S1). We initially tried to crystallize P447G with the flexible amino acid, but the glycine mutant precipitated during expression and could not be refolded, highlighting the importance of P447 for maintaining structural stability. To further understand the structural basis of their differing enzymatic activities, we compared apo-WT MKP5-CD or Cmpd 1-WT MKP5-CD with the corresponding mutants, with or without inhibitor: apo-I445A, Cmpd 1-S446, and Cmpd 1-P447V.

Superimposition of I445A onto the apo-WT structure revealed significant conformational changes across various regions of the protein (Fig. 4*A*). The structural perturbations propagate from the mutation site throughout the α4-α5 loop and β5-α3 regions. In apo-WT MKP5-CD, the loop structure is stabilized by two hydrogen bonds between residues S446 and V412, and N448 and S413. Both hydrogen-bond interactions are disrupted in the I445A mutant, with the S446-V412 and N448-S413 distances increasing to 4.57 Å and 4.70 Å, respectively, indicating loss of these stabilizing contacts and an alteration in the local loop conformation (Fig. 4*B*, S10). Additionally, the isoleucine-to-alanine substitution eliminates the buried side chain atoms Cγ1, Cγ2, and Cδ1, creating an internal cavity that destabilizes the mutant protein and contributes to reduced catalytic activity (17). This internal cavity formation, with the weakened hydrogen bond between the carbonyl oxygen of R442 and the amide nitrogen of A445, measured at 3.5 Å instead of 2.9 Å in apo-WT MKP5-CD, also disrupts the dynamics of the C-terminal end of the α3 helix and the β5-α3 loop. These changes destabilized the catalytic pocket and altered the dynamics of the α3 helix, resulting in a conformation that closely resembled the Cmpd 1-bound structure of the WT protein (Fig 4*A, C*). These structural perturbations, involving the loss of stabilizing hydrogen bonds, the formation of an internal cavity, and conformational changes in helix and loop dynamics, collectively account for the reduction in catalytic activity observed in the I445A variant.

**Figure 4.**
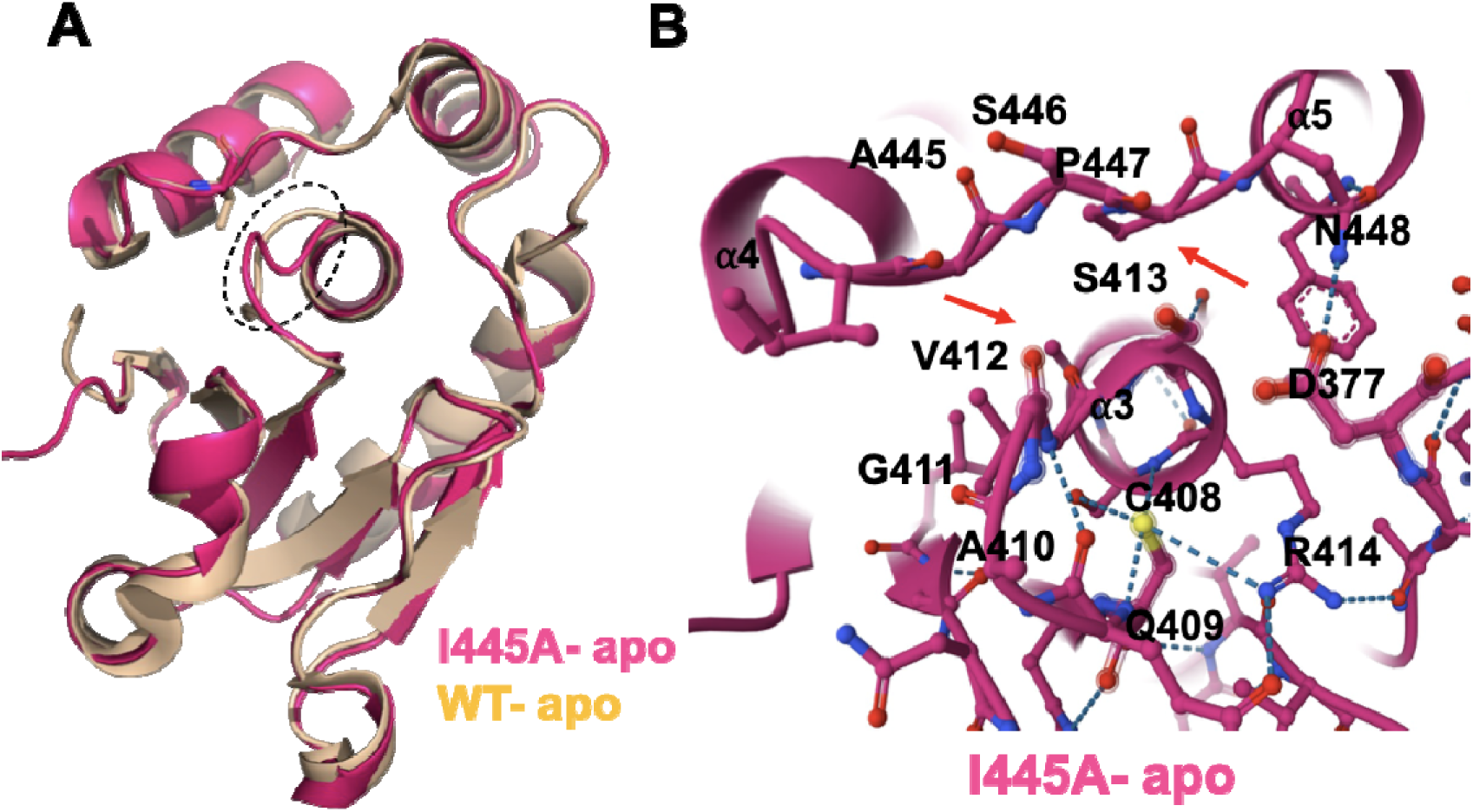
Structural comparison of WT and I445A MKP5-CD. *A*, Superposition of apo WT (wheat) and apo I445A (magenta) structures showing overall similarity except for the catalytic β5-α3 loop region. *B*, Corresponding hydrogen-bonding network in the catalytic pocket of apo I445A with missing hydrogen bonds (red arrows). Hydrogen bonds are shown as dashed lines, and key residues are displayed in stick representation. Panel A was generated in PyMOL and panel B in the Mol* viewer.

We generated the S446G and P447V complexes with Cmpd 1 and compared them to bound WT protein (Fig. 5*A*, *B*). Superposition revealed that neither mutation induces discernible conformational changes in the α4-α5 loop or its surrounding secondary structures, with well-defined electron density for Cmpd 1 in all mutant complexes (Fig. S11). For S446G, the glycine substitution eliminates the serine side chain entirely. Since the S446 side chain in WT is positioned on the solvent-exposed surface and does not form contacts with neighboring residues (Fig. 5*A*, 5*C*), its removal has minimal structural impact. This explains why S446G retains the highest catalytic activity among the four α4-α5 loop variants. In contrast, the crystallographic analysis of P447V revealed no significant alterations in active-site interactions (Fig. 5*D*), despite the reduced enzymatic activity and NMR line broadening observed for this mutant. This apparent discrepancy probably indicates that the P447V substitution affects the loop’s dynamic behavior and conformational sampling, which can be observed by NMR but are not evident in the static crystal structure.

**Figure 5.**
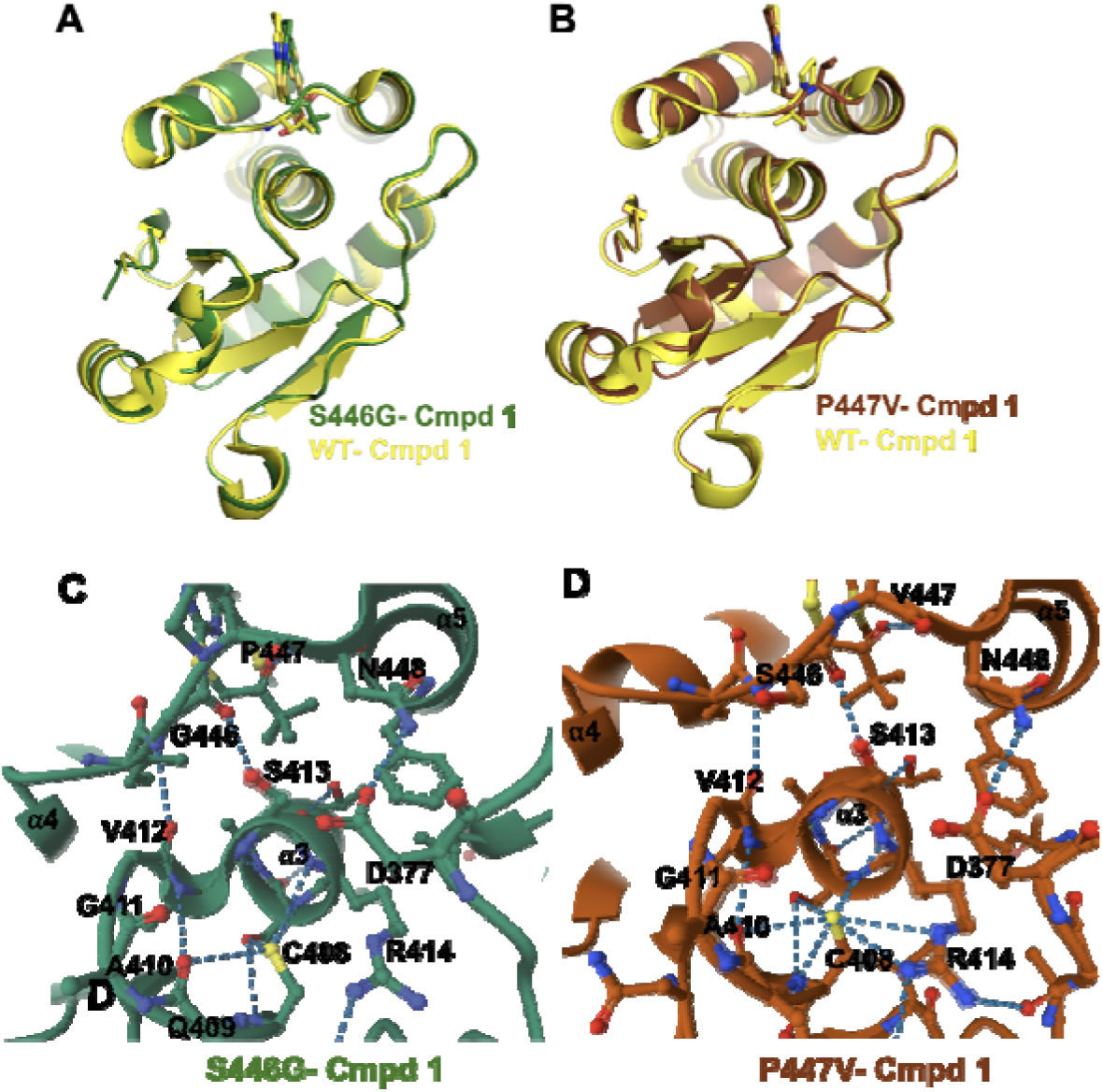
Structural comparison of Cmpd 1-bound MKP5 mutants S446G and P447V with Cmpd 1- bound WT MKP5-CD. *A*, Superposition of S446G-Cmpd 1 (green) and WT-Cmpd 1 (yellow) structures, showing high overall similarity. *B*, Superposition of P447V-Cmpd 1 (brown) and WT- Cmpd 1 (yellow) structures, which are also globally similar. *C*, Close-up views of the active-site hydrogen-bonding network **of** S446G-Cmpd 1. *D*, The same view of the active site pocket for P447V–Cmpd 1. Hydrogen bonds are shown as dashed lines, and key residues are displayed in stick representation. The figures were generated using the PyMol and Mol* molecular visualization tools. Figures *A* and *B* were generated using PyMOL, whereas panels *C* and *D* were generated using the Mol* viewer.

Given dynamic changes in the crystal structures of the variants within the α4-α5 loop that influence the structural properties of D377, Q409, S413, and R442, we mutated these residues to alanine to assess their effects on MKP5 activity. As expected, D377A, a residue essential for activity, abolished catalysis (Fig. 6*A*, *B*). When Cmpd 1 binds to MKP5, Q409 establishes new hydrogen bonding with G411 and V412. We observed that Q409A maintained ∼50% activity, indicating that this residue contributes to MKP5 catalysis with pNPP. However, this dependency was not apparent when DiFMUP was used as the substrate (Fig. 6*A, B*). Residues S413 and R442 also form new hydrogen bonds when Cmpd 1 binds, with S413A and R442A variants displaying markedly reduced MKP5 activity in assays with pNPP and DiFMUP (Fig. 6*A*, *B*). Similarly, when measured against the p38α-MAPK phosphopeptide analog, Q409A exhibited impaired catalytic activity, and both variants, S413A and R442A, completely inhibited catalysis (Fig. 6*C*).

**Figure 6.**
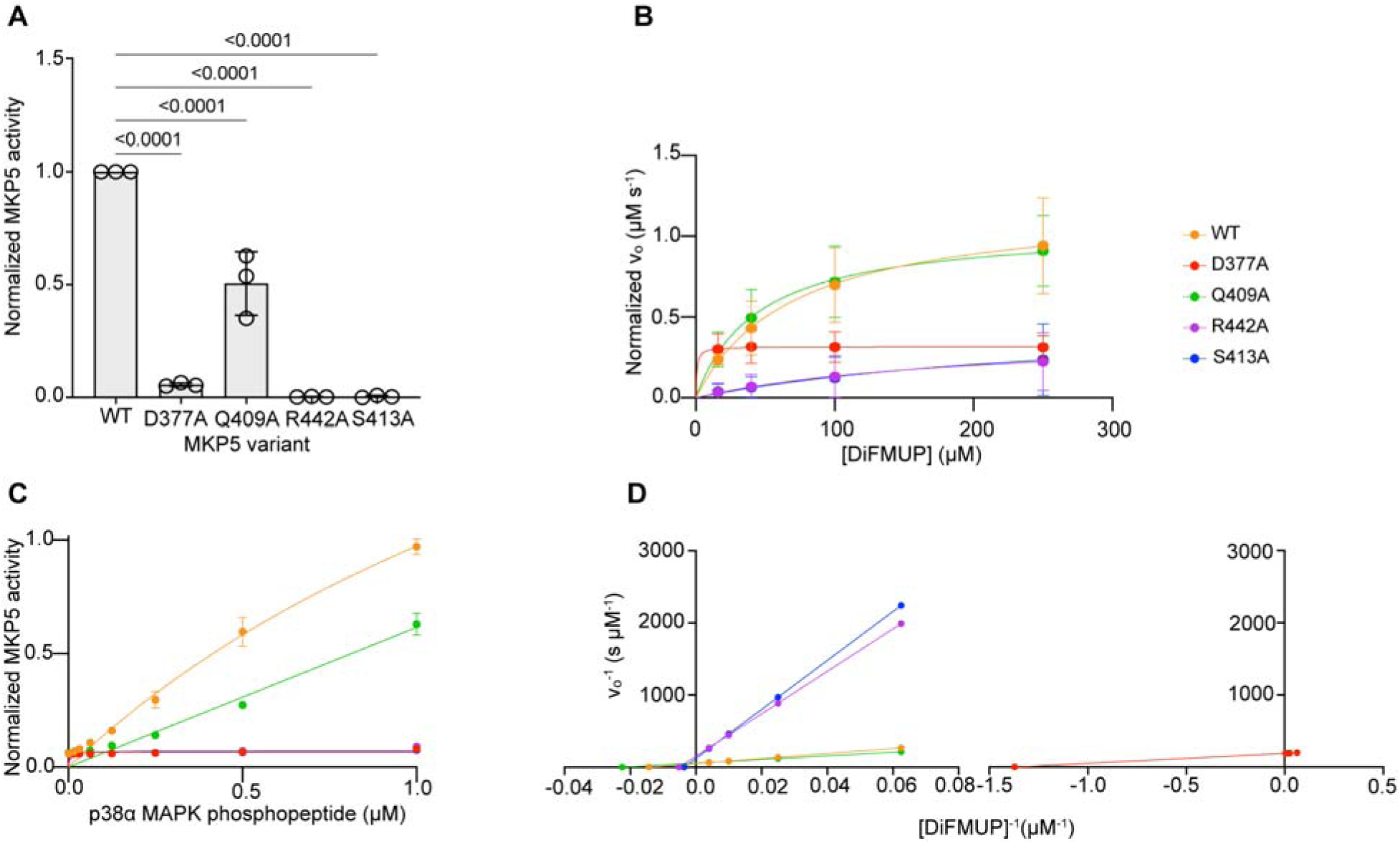
Effect of catalytic residues influenced by α4-α5 loop on MKP5 phosphatase activity. MKP5 WT, D377A, Q409A, S413A or R442A protein was assayed using *A*, *p*NPP (1.25 mM) for 10 min at 37°C. Absorbance was measured at 405 nm. Significance was determined by one-way ANOVA. *B*, DiFMUP (200 nM) for 30 min at room temperature. Fluorescence was monitored at an excitation of 358 nm and emission of 450 nm. *C*, p38 MAPK phosphopeptide for 20 min at 30°C. Malachite green was added and incubated at room temperature for 15 min. Absorbance was measured at 620 nm. *D,* Lineweaver-Burk analysis of the DiFMUP kinetic assay in *B*.

Finally, we determined the Michaelis-Menten kinetics of the variants D377A, Q409A, S413A, and R442A by measuring their initial velocities of phosphatase activity using DiFMUP (Fig. 6*D*, Table 2). Consistent with the complete lack of activity, both S413A and R442A variants exhibited a 2.8- and 4.0-fold increase in *K*_m_, respectively, as compared with WT (Fig. 6*D*, Table 2). As expected, D377A had a substantially lower *K*_m_ than WT but a 3.8-fold lower *V*_max_, indicating compromised substrate turnover. Q409A showed a lower *K*_m_ than WT but a comparable *V*_max_, consistent with the observed ∼50% retention of catalytic activity relative to WT under steady-state conditions (Fig. 6*D*, Table 2). Collectively, these data demonstrate that residues comprising the α4-α5 loop are involved in the reorganization of essential catalytic site residues upon engagement of the allosteric site. The position of the α4-α5 loop between the allosteric and catalytic sites, along with its significant conformational change upon binding Cmpd 1, indicates a role in mediating communication between these sites using residues in β4-α2, β5-α3, and α4-α5.

**Table 2:** MKP5 D377A, Q409A, S413A, and R442A kinetic parameters against DiFMUP.

|  | WT | D377A | Q409A | R442A | S413A |
| --- | --- | --- | --- | --- | --- |
| $k_{cat}$ (s<sup>-1</sup>) | 0.200 | 0.053 | 0.176 | 0.066 | 0.083 |
| $K_m$ (μM) | 69.84 | 0.7292 | 44.41 | 194.3 | 280.2 |
| $V_{max}$ (μM.s<sup>-1</sup>) | 0.020 | 0.005 | 0.018 | 0.007 | 0.008 |
| $k_{cat}/K_m$ (μM<sup>-1</sup>.s<sup>-1</sup>) | 2.86 x 10<sup>-3</sup> | 7.26 x 10<sup>-2</sup> | 3.97 x 10<sup>-3</sup> | 3.40 x 10<sup>-4</sup> | 2.94 x 10<sup>-4</sup> |

## Discussion

The MAPKs are specifically activated by dual phosphorylation of the T-x-Y motif that resides in the activation loop (11). Regulation of MAPK activity is achieved through the integrated actions of the MAPK kinases that phosphorylate, and the opposing actions of the MKPs that dephosphorylate the T-X-Y motif. Much is known about the regulation of the MAPK kinases and MAPK phosphorylation (10,11). In contrast, a similarly comprehensive understanding of how the MKPs are regulated to mediate MAPK dephosphorylation has yet to be fully realized. There are 10 active MKPs, all of which exhibit highly selective substrate preference to either ERK1/2, p38 MAPK, or JNK (18). The current view is that MKP activation occurs through interaction of its kinase-binding domain (KBD) with the MAPK substrate. A number of MKPs, notably MKP3, have been shown to be regulated in this manner (19). Although not as robust as the activation of MKP3 upon ERK1/2 binding, other MKPs, such as MKP5 have been reported to be activated by p38 MAPK and JNK binding (20). Structurally, the MKP5 CD favors an activated state and differs from that of the MKP3 CD (21). However, the identification of the allosteric site on MKP5 suggests that there are additional molecular mechanisms that exist which further lead to its activation upon p38 MAPK and JNK binding.

Previously, we identified a small-molecule allosteric inhibitor for MKP5 that bound to a region distal to the catalytic site, which we showed to be critical for activity (16). Furthermore, the site bound by the allosteric inhibitor also constituted a location on MKP5 that is required for the binding of both p38 MAPK and JNK (15). These observations prompted us to posit that engagement of p38 MAPK and JNK at the allosteric site induces conformational modifications within the PTP domain that promote catalysis. The residue Y435 was required for p38 MAPK/JNK and MKP5 allosteric inhibitor binding, indicating that its engagement serves as a platform for transmitting conformational alterations to the critical C408 catalytic residue (15). Although it is known that the KBD facilitates MKP5 binding to p38 MAPK/JNK (22,23), identification of the allosteric site indicates that additional regulatory mechanisms of these interactions are operative for the control of catalytic activity. In this regard, conformational changes in the α4-α5 loop play an important regulatory role on the effects of p38 MAPK and JNK binding and impact the integrity of the catalytic pocket.

Upon Cmpd 1 binding to WT MKP5-CD, there is a notable movement of P447 in the α4-α5 loop and a shift of β5-α3 residues. Our initial focus was on the conformational change in P447, but we knew that its distinctive cyclic backbone, rigid structure, and secondary amine would make it difficult for any mutant to elucidate its role. The generation of a Gly mutant was insoluble, but the P447V mutation was crystallized in the Cmpd 1-bound state, exhibited weak enzymatic activity in the apo-state, showed NMR resonance line broadening, and offered preliminary insight into the conformational change of P447. However, we identified mutations at other positions adjacent to P447 and found the N448A mutation in both the apo- and Cmpd 1-bound forms, providing an excellent understanding of the α4-α5 loop in general. N448A was inactive because of dysfunctional interactions with the important D377 amino acid, and induced changes in β5-α3, β4-α2 (only in apo), and β3-α5 and the end of the helix α3 that affect catalysis. These findings provide critical insight into the enzyme’s structure-function relationship and highlight how non-catalytic residues can be indispensable for maintaining enzymatic competence.

Despite MKP5’s high basal activity (23), p38 MAPK and JNK binding still increase its activity (20). Therefore, it is conceivable that the α4-α5 loop also participates in promoting MKP5 activity upon p38 MAPK and JNK binding. It is also plausible that small-molecule inhibitors targeting the α4-α5 loop could be effective antagonists of MKP5 by either disrupting or locking the α4-α5 loop in a less favorable conformation. Indeed, in silico modeling of MKP4 inhibitors has suggested that the α4-α5 loop could be targeted to achieve enzyme inhibition (24). The α4-α5 loop, which forms part of the catalytic cleft, is conserved amongst several other MKPs (e.g., MKP4, PAC-1, PYST2, MKP7, and MKP3), suggesting the possibility of a conserved mechanism of activation occurring through the α4-α5 loop amongst the MKPs. Indeed, MKP7 has an allosteric pocket that also binds p38 MAPK and JNK and possesses a conserved α4-α5 loop (25). Mutation of the allosteric pocket on MKP7, similarly inhibits its activity (25).

Given the emergence of MKP5 as a potential therapeutic target for diseases such as idiopathic pulmonary fibrosis (26), cardiac fibrosis (27), and dystrophic muscle disease (28), the rationale for developing small-molecule MKP5 inhibitors is compelling. Development of allosteric MKP inhibitors extends beyond MKP5, as other MKPs such as MKP1 (*DUSP1*) and MKP3 (*DUSP6*) have been heavily implicated in cancer drug resistance and targeting of these MKPs could provide enormous therapeutic benefit for the treatment of certain cancers. Finally, the implications of this study support a two-step mode of MKP activation on a global level. MAPK binding to the KBD could occur first, bringing the MAPK in to close proximity of the PTP domain followed by a second step involving engagement at the allosteric site, which then induces the conformational changes through the α4-α5 loop to the active site. This is consistent with a ligand-induced activation model and would not negate the necessity of the KBD. Importantly, small-molecule allosteric inhibitors could disrupt the interaction of incoming MAPK substrates, providing an additional mechanism of MKP inhibition.

## Experimental Procedures

### Protein expression and purification

Given the large conformational change observed in wild-type (WT) MKP5 catalytic domain (CD), we used Ala/Gly/Val mutagenesis (16). For P447, we used Val due to insolubility of the Gly mutants. The WT and variants (I445A, S446G, P447V, N448A, S413A, D377A and R442A) of the MKP5 CD were cloned and expressed as previously described (29). Mutant constructs were engineered using Pfu Turbo DNA polymerase (Agilent). Briefly, MKP5-CD cDNA mutants were cloned into a pET28a vector containing a TEV cleavage site followed by transformation into BL21 Gold (DE3) cells. The resultant mutagenized vectors were confirmed by DNA sequencing.

Mutant MKP5-CD clones were induced using 1 mM IPTG and expressed overnight at 20°C. Following induction, the cells were suspended in lysis buffer containing 20 mM HEPES (pH 7.4), 500 mM NaCl, 10% glycerol, 5 mM imidazole, 2 mM β-mercaptoethanol, DNase 1, and complete EDTA-free protease inhibitor. Subsequently, the soluble protein fraction was loaded onto a pre-packed Ni-NTA column to purify the His-tagged protein and the tag was cleaved using TEV protease. The catalytic mutants were subjected to gel filtration chromatography using a Hi-Load 16/600 Superdex75 GL column to obtain pure, monodispersed protein samples. Following concentration, the protein was stored in 20 mM Tris, pH 8.0, 150 mM NaCl, 5% glycerol and 5 mM dithiothreitol at −80°C.

### Phosphatase activity assays

For MKP5 phosphatase activity assays using *p*-nitrophenol phosphate (*p*NPP) as substrate, MKP5 WT or MKP5 variant proteins were diluted to a working concentration and the reaction was initiated with 40 µL of *p*NPP reaction buffer. The reaction buffer consisted of a final concentration of 20 mM HEPES pH 7.4, 120 mM NaCl, 5 mM DTT, and 12.5 mM *p*NPP, with the final concentration of MKP5 WT or MKP5 variant proteins (2.5 µM). Samples were run in triplicate in a 96-well plate. The reaction was incubated for 10 minutes at 37°C. The reaction was quenched by adding 200 µL of 0.2 N NaOH, and absorbance was measured using an Agilent Biotek Synergy H1 plate reader at 405nm. Absorbance was converted to turnover using an extinction coefficient of 17,800 M^-1^ cm^-1^. For DiFMUP (6,8-Difluoro-4-methylumbelliferyl phosphate) assays, DiFMUP was purchased from Thermo Fisher Scientific (Lot no. 2016387). MKP5 WT or MKP5 variant proteins were diluted to a final concentration of 200 nM in 50 mM Tris-HCl, pH 8.0, 5 mM DTT, and 0.05% Tween-20. Final concentrations of DiFMUP were 250 µM, 100 µM, 40 µM, and 16 µM. Samples were run in triplicate on 96-well plates. To determine the kinetics, 100 nM MKP5 WT or variants were incubated in the reaction buffer and the reaction initiated by the addition of DiFMUP. Reactions proceeded for 30 min at room temperature and progress was monitored using an Agilent Biotek Synergy H1 plate reader with excitation at 358 nm and emission at 450 nm. Initial velocity was determined by linear regression. These velocities were plotted against DiFMUP concentration which was fit using a Michaelis-Menten curve. p38 MAPK phosphopeptide analog (Asp-Asp-Glu-Nle-pThr-Gly-pTyr-Val-Ala-Thr-Arg) and MKP5 WT or variant proteins were incubated in a 96-well plate for 20 minutes at 30°C in PTP buffer, which consisted of 50 mM Tris-HCl (pH 7.2) and 1 mM EDTA. The final concentration of MKP5 WT or mutant protein was 1 μM, and the final concentration of p38 MAPK phosphopeptide was 1 mM, 0.5 mM, 0.25 mM, 0.125 mM, 0.0625 mM, 0.03125 mM, and 0.015625 mM. 200 μL of malachite green development reagent was added and incubated for 15 minutes at room temperature. The malachite green development reagent consisted of 1.6 N HCl, 0.027% malachite green dye, and 1.68% ammonium molybdate. Absorbance was measured at 620 nm using an Agilent Biotek Synergy H1 plate reader.

### NMR spectroscopy

MKP5 NMR samples were expressed in M9 minimal media with ^15^NH_4_Cl and ^13^C_6_H_12_O_6_ (Cambridge Isotope Laboratories) as the sole nitrogen and carbon sources, respectively. The cells were cultured at 37°C to an OD_600_ of 0.8 – 1.0, induced with 1 mM IPTG, and then incubated at 20°C for an additional 16 to 18 hours. Isotopically labeled MKP5-CD WT, I445A, S446G, P447V, and N448A samples were purified as described earlier. The samples were dialyzed into a buffer containing 40 mM sodium phosphate, 2 mM DTT, and 1 mM EDTA at pH 7.4. Samples were concentrated to 0.5 mM and 7.5% D_2_O (v/v) was added prior to NMR experiments. All NMR experiments were collected on a Bruker Avance NEO 600 MHz spectrometer at 25°C equipped with pulsed field gradients and a triple-resonance cryoprobe. NMR spectra were processed with NMRPipe and analyzed in Sparky. The backbone resonance assignments of MKP5-CD were obtained from our prior work (BMRB accession code #53231) and transferred to MKP5 variants. ^1^H^15^N combined chemical shift perturbations (Δδ) were determined from ^1^H^15^N TROSY HSQC spectra by 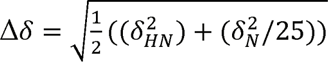, where δ_HN_ and δ_N=_ δ*_WT_* - δ*_Variant_* |. Significant chemical shift perturbations were determined as values 1.5σ above the 10% trimmed mean of all data sets.

### CD Spectroscopy

MKP5-CD samples (10 μM) were dialyzed into a buffer containing 40 mM sodium phosphate and 1 mM EDTA at pH 7.4. Thermal unfolding CD experiments were collected on a JASCO J-815 spectropolarimeter equipped with a variable-temperature Peltier device using a 2 mm quartz cuvette. Experiments were collected on apo MKP5-CD WT and variant samples for 1 hour at room temperature prior to data collection. Denaturation curves were recorded at 223 nm over a temperature range of 20 to 90°C. *T_m_* values were determined via nonlinear curve fitting in GraphPad Prism.

### Crystallization and structure determination

The MKP5 CD variants I445A, S446G, P447V, and N448A were attempted to crystallize with and without Compound 1 (Cmpd 1; 3,3-dimethyl-1-((9-(methylthio)-5,6-dihydrothieno[3,4-h]quinazolin-2-yl)thio)butan-2-one)(16) using the sitting-drop method. In a 96-well plate, drops of 300 nl protein mixed with 200 nl reservoir solution were set using the Formulatrix NT8 drop setter. Crystals of the S446G, P447V, and N448A mutants in complex with Cmpd 1 were obtained under the following conditions: 0.2 M Tricine pH 7.4 with sodium hydroxide, 20% (w/v) PEG 4000; 0.2 M N-(2-Acetamido)iminodiacetic acid (pH 6.7), 20% (w/v) PEG 4000; and 0.1 M Tris pH 6.5 with 2 M ammonium sulfate, respectively. The apo crystals of I445A and N448A were grown in 0.1 M imidazole, pH 8.0, 2.5 M sodium chloride and 0.1M Tris pH6.5, 2.0M Ammonium Sulfate respectively.

X-ray crystallographic data sets were collected at the Brookhaven Laboratory National Synchrotron Light Source II, at the NYX, AMX and FMX beamlines. Molecular replacement was performed using Phaser, with the MKP5-CD structure in complex with Cmpd 1 (PDB: 6MC1) as the search model. The 2|Fo| - |Fc| and |Fo| - |Fc| Fourier maps were analyzed, revealing electron density for the Cmpd 1. Structure refinement was performed using Phenix.refine with multiple rounds of refinement cycles and model building in Coot (30) to achieve the final structure. Stereochemistry assessment with MOLPROBITY confirmed good quality in the crystal structures (31). The polder maps were calculated in Phenix to validate ligand density (32). Data collection, scaling, and refinement statistics are summarized in Table S1. The structural coordinates of the apo and holo complex have been deposited in the RCSB Protein Data Bank with the following codes: S446G-Cmpd 1 (9NSB.pdb), P447V-Cmpd 1 (9OK9.pdb), N448A-Cmpd 1 (9Q7X.pdb), apo I445A (9Y55.pdb), and apo N448A (9NYM.pdb). Electron density of the variants to S446G, P447V and N448A variants is shown in Fig. S11.

## Data Availability

All data supporting the results of this study are available in the article, including the Supplementary Information and source data. MKP5 structures have been deposited to PDB under accession numbers 9NSB (S446G-Cmpd 1), 9OK9 (P447V-Cmpd 1), 9Q7X (N448A-Cmpd 1, 9Y55 (apo I445A), and 9NYM (apo N448A). NMR chemical shift and peak height data generated in this study are provided in the Supplementary Information.

## Supporting Information

This article contains supporting information.

## Author Contributions

R.M., L.G., and L.B.; Expressed and purified MKP5 proteins, R.M.; performed X-ray crystallography studies and associated data analysis, L.G.; performed all enzymatic experiments and associated data analysis, E.S.; performed NMR, CD experiments and associated data analysis; G.P.L., supervised NMR spectroscopy, E.J.L.; conceived the study and supervised X-ray crystallography; A.M.B.: conceived the study and supervised enzymatic studies. The manuscript was written through contributions of R.M., L.G., G.P.L., E.J.L., and A.M.B.

## Funding and Additional Information

This work was partially supported by NIH grants R01 GM144451 to G.P.L. and R01 HL158876 to A.M.B. and E.L. A.M.B. was also partially supported by R01 AR080152. L.G. was supported by T32 GM149444.

