## Supplemental Data for "Functional coupling between the α4 – α5 loop and allosteric site on MKP5 is a critical determinant of catalysis"

*; The University of Tennessee Health Science Center, Department of Pharmaceutical Sciences, Memphis, Tennessee, USA

**SUPPLEMENTAL INFORMATION**

**Table S1.** Data collection and refinement statistics.

**Figure S1.** Secondary structure representation for the catalytic domain of MKP5.

**Figure S2**. Structural alignment of WT-apo and Cmpd 1-bound complex.

**Figure S3 – S6.** NMR spectra of WT MKP5-CD and α4 – α5 loop variants.

**Figure S7.** Thermal stability of WT MKP5-CD and α4 – α5 loop variants.

**Figure S8**. Comparison of rotations and distances of D377 side chains in Cmpd 1-WT MKP5 CD and Cmpd 1-N448 in the crystal structures.

**Figure S9**. Orientations of the D377 and N488A are shown for the N448A mutation structures, and the wild-type 2.7 Å hydrogen-bond interaction between these residues.

**Figure S10.** Depiction of altered α3 helix dynamics in the I445A-apo structure.

**Figure S11**. Electron density of Cmpd 1 binding to S446G, P447V, and N448A variants.

**Table S1. Data collection and refinement statistics.**

|  | **S446G-Cmpd 1**  (9NSB.pdb) | **P447V-Cmpd 1**  (9OK9.pdb) | **N448A-Cmpd 1**  (9Q7X.pdb) | **N448A-Apo**  (9NYM.pdb) | **I445A-apo**  (9Y55.pdb) |
| --- | --- | --- | --- | --- | --- |
| **Wavelength** | 0.979 | 0.979 | 0.978 | 0.978 | 0.979 |
| **Resolution range** | 34.5  - 2.5 (2.589  - 2.5) | 34.25 - 3.0 (3.107 - 3.0) | 51.39 - 2.95 (3.055 - 2.95) | 34.46  - 2.0 (2.071  - 2.0) | 166.2 - 3.502 (3.627 - 3.502) |
| **Unit Cell** | 95.697 99.559 135.301 90.00 90.00 90.00 | 92.74 98.25 136.99 90.00 90.00 90.00 | 65.86 129.95 84.0105 90 91.38 90 | 39.3101 40.5305 57.3797 74.6199 78.86 61.55 | 106.404 110.048 166.234 90 90.205 90 |
| **Space group** | P 21 21 21 | P 21 21 21 | P 1 21 1 | P 1 | P 1 21 1 |
| **Unique reflections** | 45398 (4478) | 23966 (4089) | 54469 (5725) | 18191 (1464) | 46662 (4844) |
| **Multiplicity** | 13.1 (12.7) | 3.7 (3.8) | 1.9 (1.9) | 1.7 (1.8) | 2.0 (2.0) |
| **Completeness (%)** | 99.90 (99.89) | 93.03 (99.76) | 95.05 (99.56) | 89.90 (95.7) | 95.89 (99.90) |
| **Mean I/sigma(I)** | 11.89 (1.14) | 10.0 (1.5) | 8.67 (1.83) | 5.0 (2.7) | 6.64 (1.55) |
| **R-merge** | 0.1714 (2.021) | 0.094(1.030) | 0.09734 (0.4811) | 0.061 (0.144) | 0.1071 (0.5164) |
| **CC1/2** | 0.998 (0.641) | 0.998(0.620) | 0.973 (0.55) | 0.981 (0.944) | 0.991 (0.437) |
| **Reflections used in refinement** | 45378 (4473) | 23921 (2515) | 28357 (2943) | 18115 (1891) | 46626 (4844) |
| **Reflections used for R-free** | 2368 (239) | 1168 (117) | 1480 (163) | 882 (103) | 2176 (199) |
| **R-work** | 0.2189 (0.3682) | 0.1969 (0.3542) | 0.1888 (0.2830) | 0.2058 (0.2157) | 0.2183 (0.3166) |
| **R-free** | 0.2697 (0.4055) | 0.2402 (0.4354) | 0.2440 (0.3453) | 0.2087 (0.2116) | 0.2424 (0.3199) |
| **Number of non-hydrogen atoms** | 7101 | 7231 | 7229 | 2490 | 14177 |
| **macromolecules** | 6963 | 7093 | 7091 | 2347 | 14177 |
| **ligands** | 138 | 138 | 138 | 10 | - |
| **Protein residues** | 884 | 884 | 882 | 294 | 1764 |
| **RMS(bonds)** | 0.013 | 0.01 | 0.004 | 0.008 | 0.003 |
| **RMS(angles)** | 1.52 | 1.33 | 0.71 | 0.89 | 0.64 |
| **Ramachandran favored (%)** | 96.1 | 92.78 | 93.91 | 95.86 | 93.62 |
| **Ramachandran allowed (%)** | 3.9 | 7.22 | 5.86 | 4.14 | 5.80 |
| **Ramachandran outliers (%)** | 0 | 0 | 0.23 | 0 | 0.57 |
| **Rotamer outliers (%)** | 4.84 | 0.13 | 0.92 | 2.81 | 0.07 |
| **Clashscore** | 9.84 | 7.32 | 7.17 | 6.86 | 7.54 |
| **Average B-factor** | 68.86 | 90.75 | 49.19 | 27.81 | 96.17 |
| **Macromolecules** | 68.89 | 90.71 | 48.89 | 27.36 | 96.17 |
| **Ligands** | 67.13 | 92.73 | 64.60 | 27.88 | - |

Statistics for the highest-resolution shell are shown in parentheses

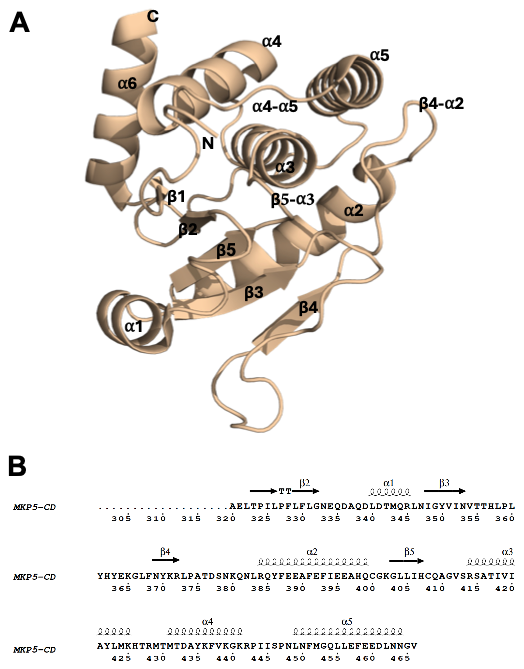

**Figure S1**. **Secondary structure representation for the catalytic domain of MKP5.** *A,* Structure of MKP5 catalytic domain (CD); *B*, MKP5 sequence is annotated with schematic representations of their corresponding secondary structural elements shown above. Panel *B* was generated using ESPript (http://espript. ibcp.fr/ESPript/ESPript/index.php). The figure was generated using the PyMol viewer.

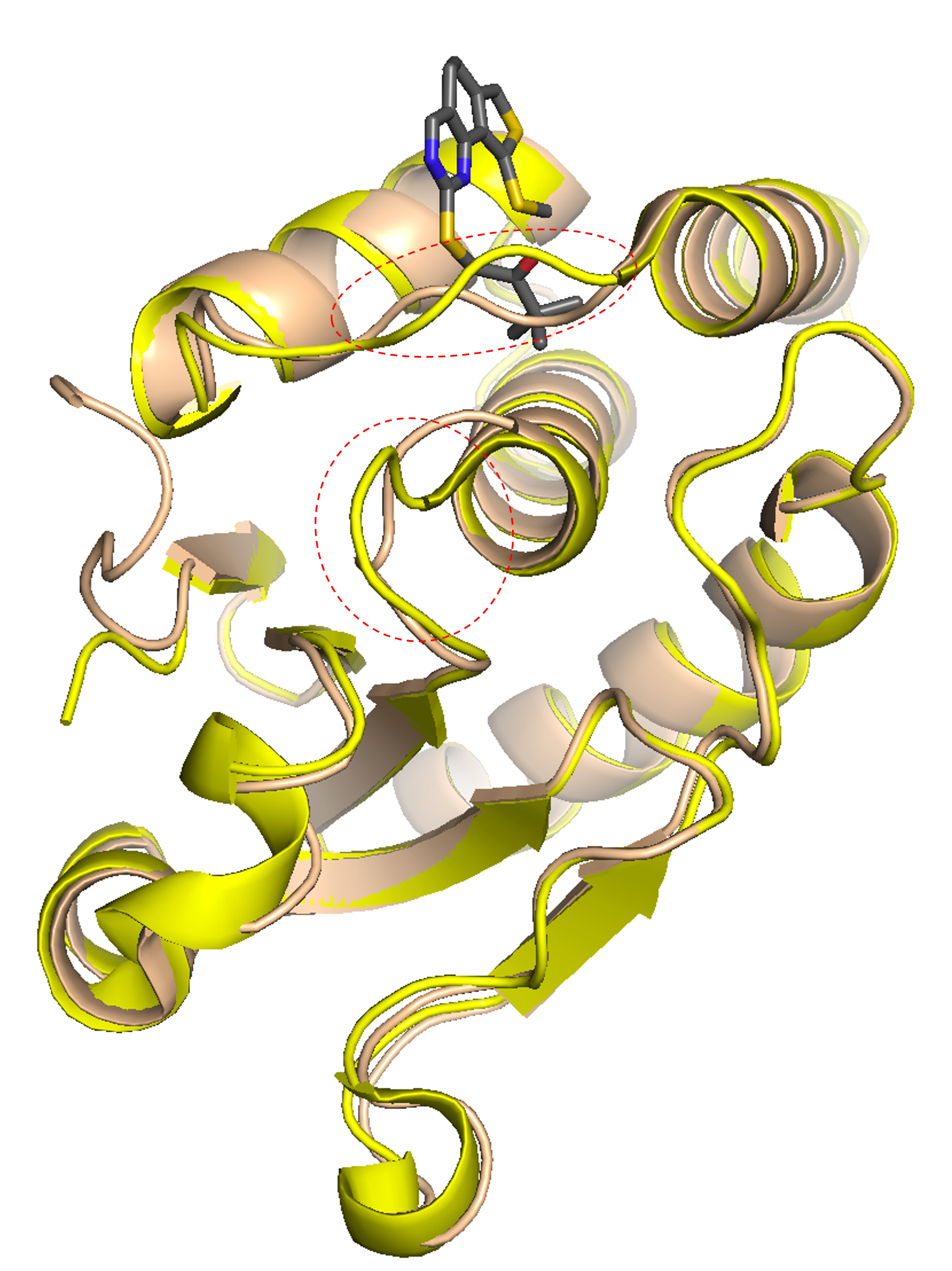

**Figure S2**. **Structural alignment of WT-apo (2OUD, Wheat) and Cmpd 1-bound MKP5 complex (6MC1, Yellow).** The WT-apo (wheat) undergoes several conformational changes upon binding of allosteric inhibitor (grey), forming the Cmpd 1-MKP5 complex (yellow) which is responsible for loss of catalytic activity. The Cmpd 1-MKP5 conformational change is observed directly below the four residues, ^445^ISPN^448^ of the α4-α5 loop. The other conformational change is the β5-α3 loop, which exhibits an intertwined conformational change between apo-WT MKP5 (wheat) and 6MC1. The figures were generated using the PyMol viewer.

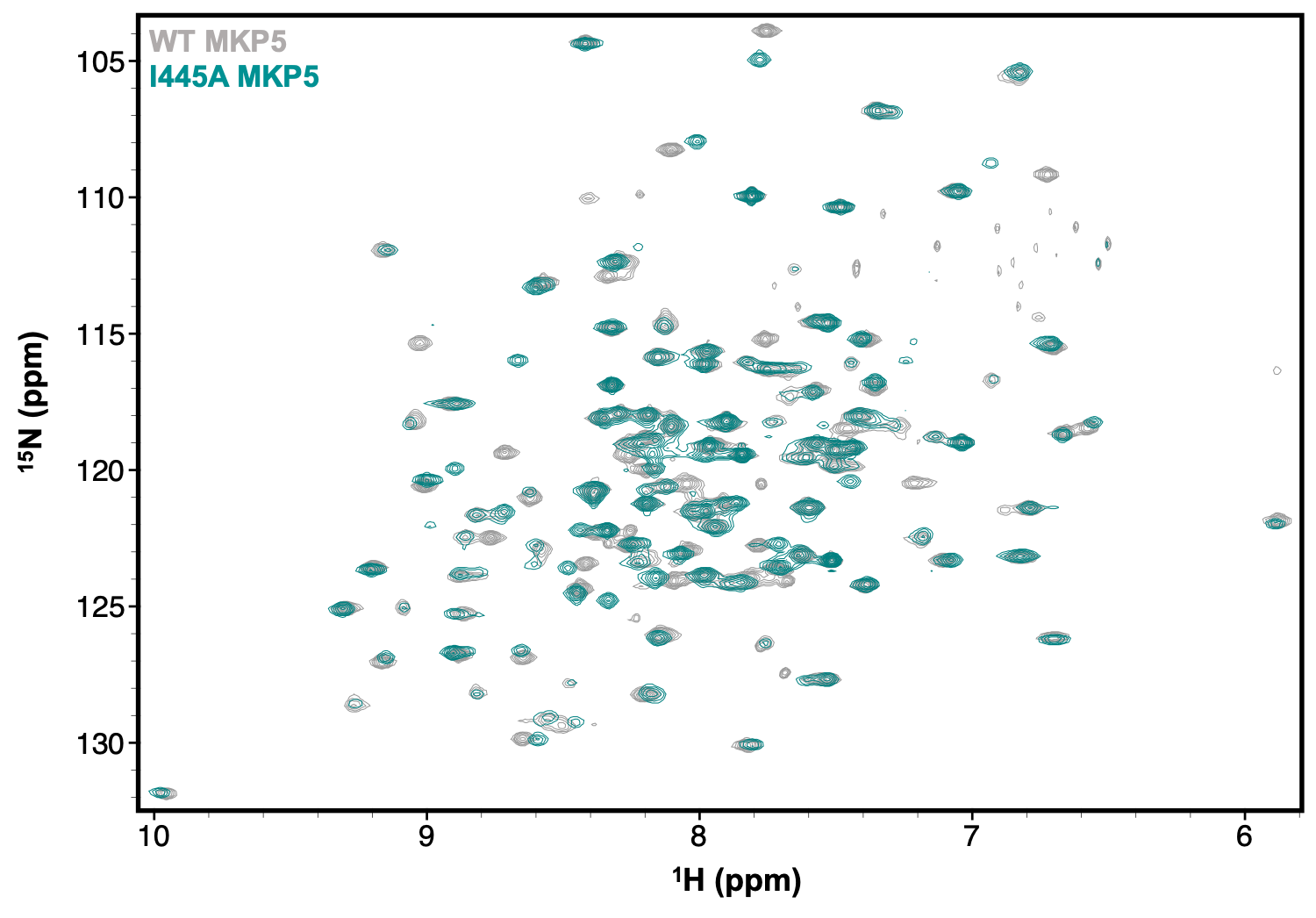

**Figure S3.** ^1^H^15^N HSQC NMR spectra of WT MKP5-CD (gray) and I445A MKP5-CD (cyan) collected at 14.1 T and 25ºC.

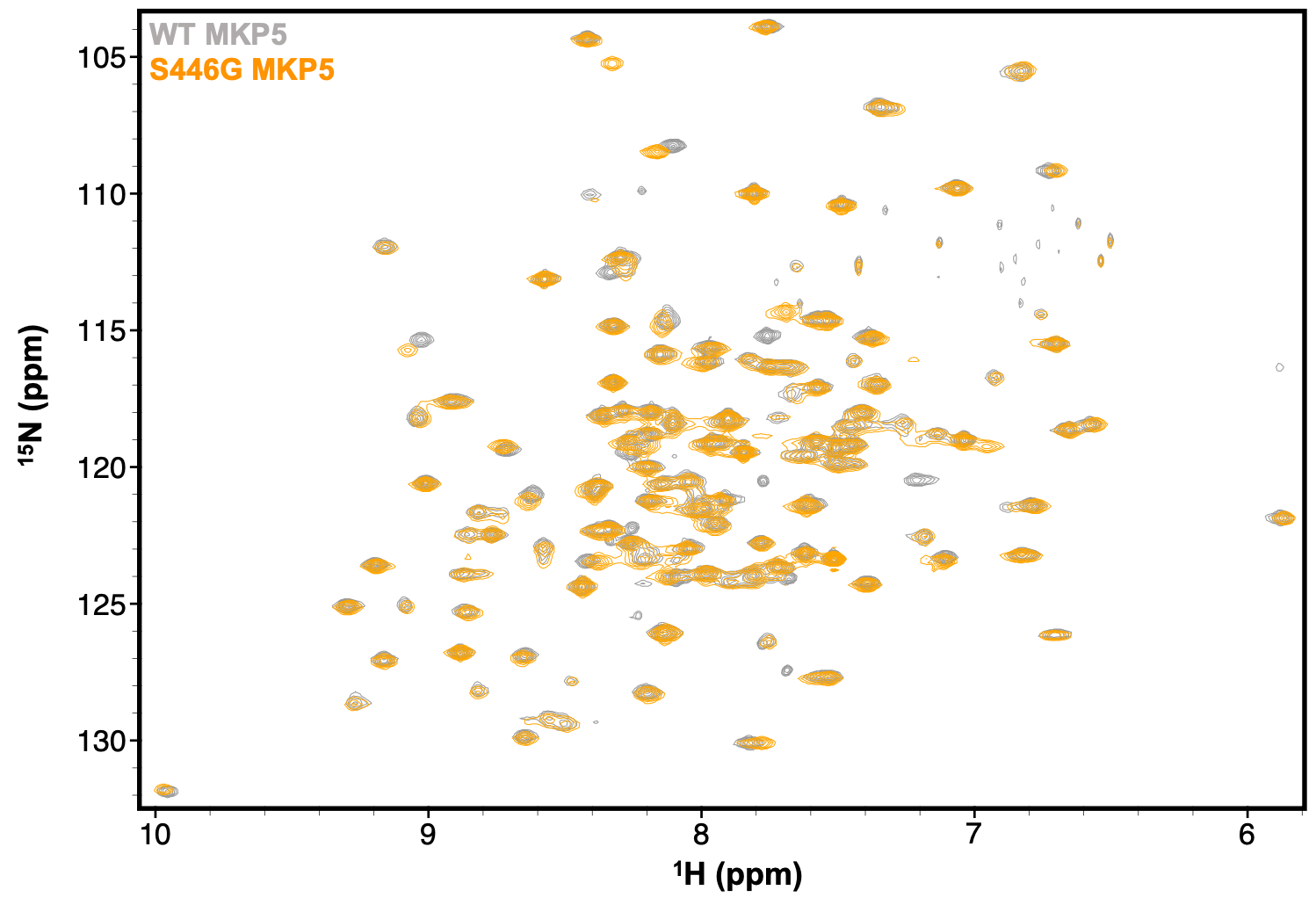

**Figure S4.** ^1^H^15^N HSQC NMR spectra of WT MKP5-CD (gray) and S446G MKP5-CD (orange) collected at 14.1 T and 25ºC

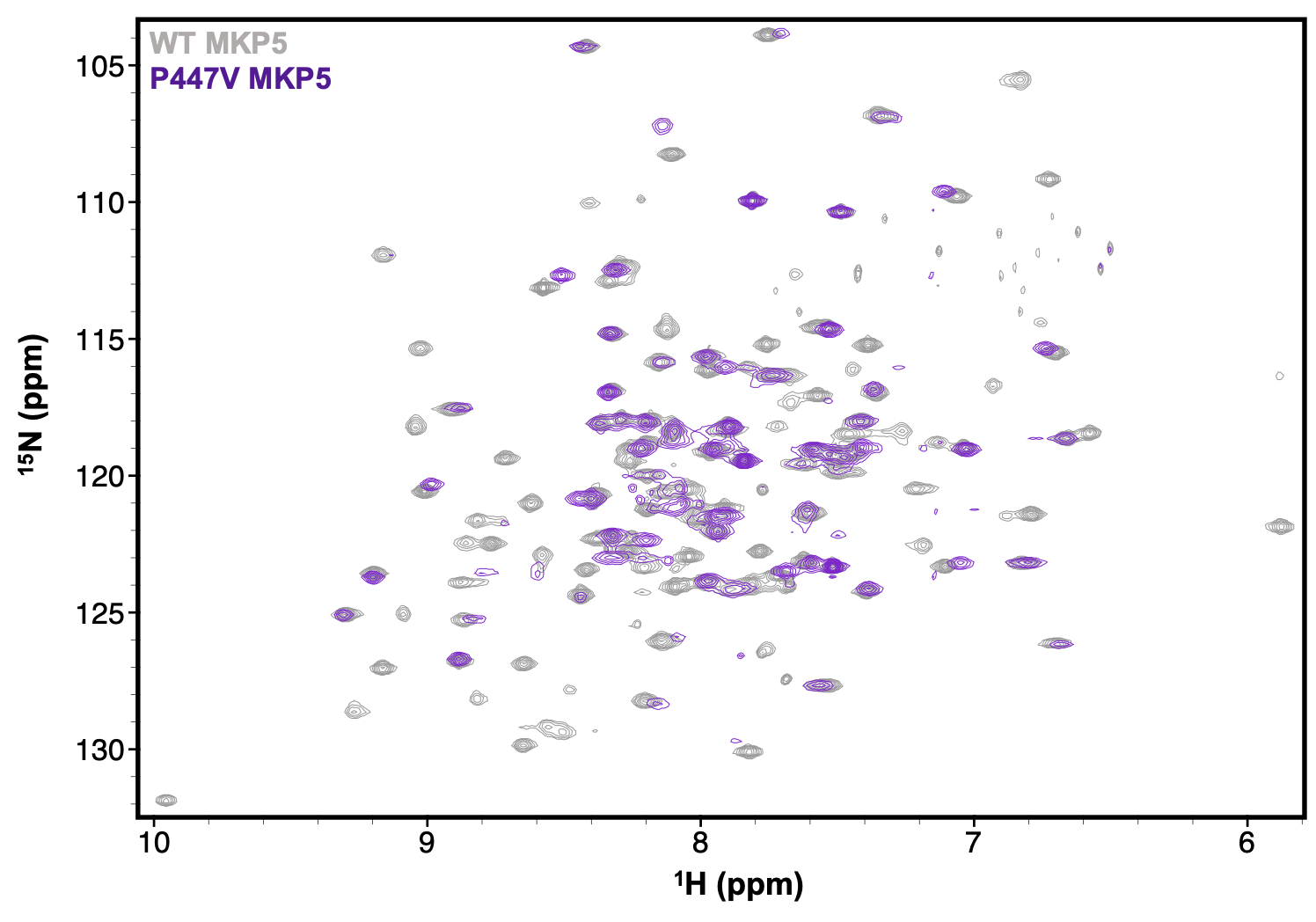

**Figure S5.** ^1^H^15^N HSQC NMR spectra of WT MKP5-CD (gray) and P447V MKP5-CD (purple) collected at 14.1 T and 25ºC

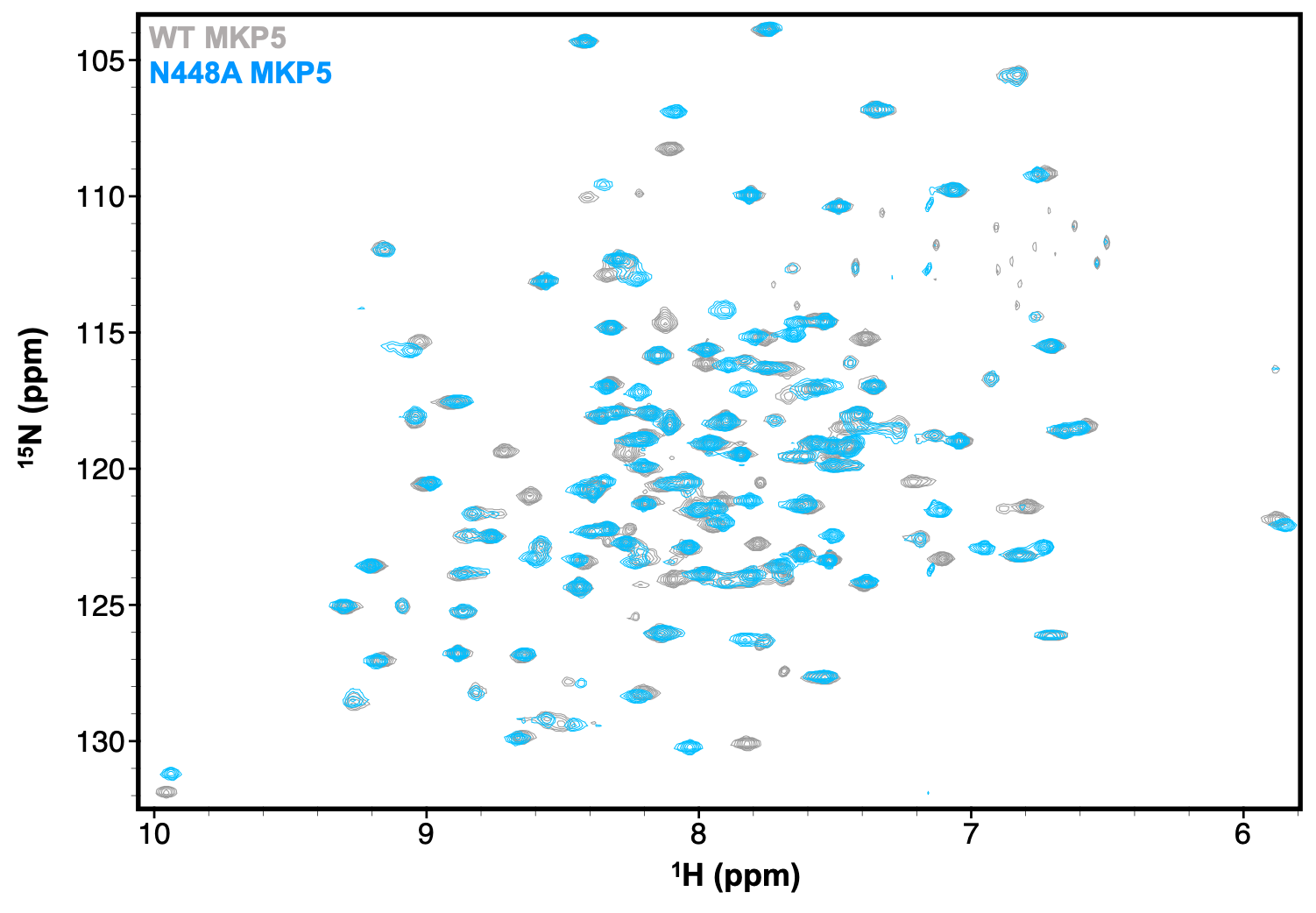

**Figure S6.** ^1^H^15^N HSQC NMR spectra of WT MKP5-CD (gray) and N448A MKP5-CD (blue) collected at 14.1 T and 25ºC

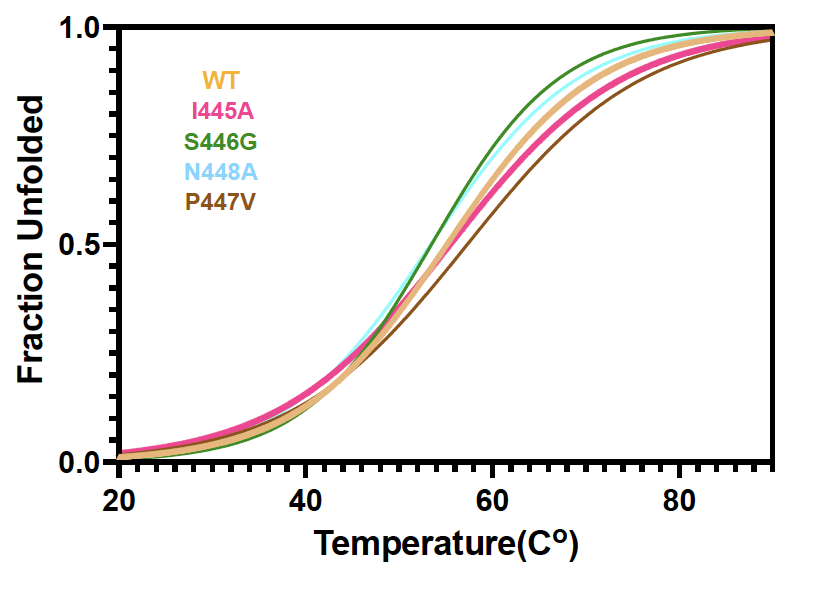

**Figure S7.** **Thermal stability of WT MKP5-CD and its loop variants.** Thermal unfolding curves derived from CD spectroscopy are shown for WT MKP5-CD (wheat), I445A (magenta), S446G (green), P447V (brown) and N448A (cyan). *T*_m_ values for WT MKP5-CD, I445A, S446G, P447V and N448A were 56.2, 57.5, 53.8, 58.4 and 53.9, respectively were determined via nonlinear curve fitting in GraphPad Prism.

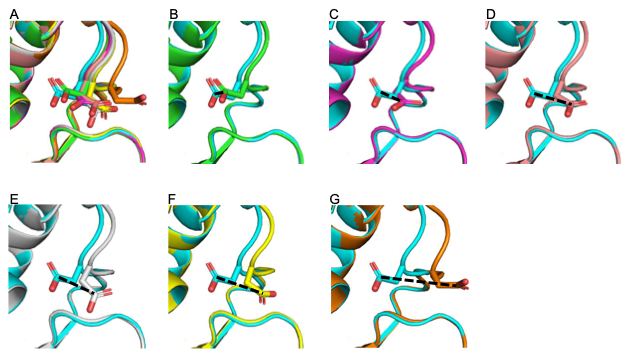

**Figure S8. Comparison of various positions for D377 from Cmpd 1-N488A MKP5 CD relative to Cmpd 1-WT MKP5 CD.** Cmpd 1-N448A MKP5 CD and Cmpd 1-WT MKP5 CD include six independent MKP5 catalytic domains in their respective crystal structures. Both proteins superimpose well overall, except near D377 in the N488A mutant. The Cg atoms illustrate the distances between the two proteins (black dashed lines). *A,* WT D377 (aqua) adopts a single orientation across all six WT MKP5 catalytic domains, while D377 of Cmpd 1-N488A exhibits six different orientations with varying rotations and distances relative to WT D377. The distances of Cg of D377 for each Cmpd 1-N448A relative to the corresponding atom in WT are: *B,* 0.7 Å; *C,* 2 Å; *D,* 3 Å; *E,* 3.3 Å; *F*, 3.8 Å, and *G*; 6.1 Å. Structures of D377 residue of Cmpd 1-N488A in *B*, *C*, *D*, and *E* show differences in Ca atoms that range from 0.5 to 0.8 Å in the β2-α4 loop compared to WT MKP5. *F* and *G* display distances in the β2-α4 loop of 1.0 Å and 2.5 Å, respectively, relative to the WT.

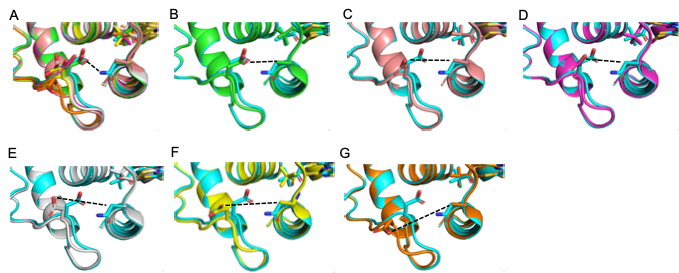

**Figure S9. Orientation of D377 and N488 in WT and mutant structures.** (A) The 2.7 Å hydrogen bond between WT D377 and N488 is illustrated (dashed black line). All D377 residues from N488A are overlaid on the WT MKP5 catalytic domain. The distances between D377 and N448A are shown in dashed lines: (B) 4.7 Å, (C) 7.2 Å, (D) 8.1 Å, (E) 9.1Å, (F) 9.2 Å, and (G) 11.0 Å.

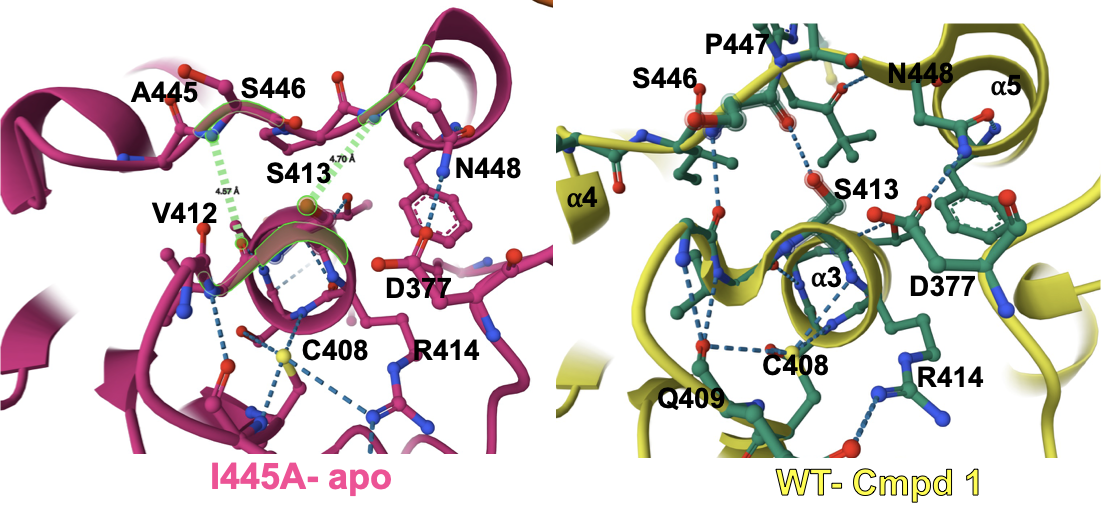

**Figure S10**. **Depiction of altered α3 helix and β5-α3 dynamics in the I445A-apo structure.** The left and right figures represent the active-site hydrogen-bonding network of apo I445A (pink) and Cmpd 1-WT (yellow). Key interactions are shown as dashed lines. The distances from the amide group of S446 to the hydroxyl group of V412, and from N448 to S413, are 4.57 Å and 4.70 Å, respectively. These measurements lead to changes at the α3 helix and the β5-α3 loop relative to WT-Cmpd 1, resembling the apo-WT structure, with the notable exception of the conformation change of α4-α5 due to the absence of Cmpd 1. The figures were generated using the Mol* molecular visualization tool.

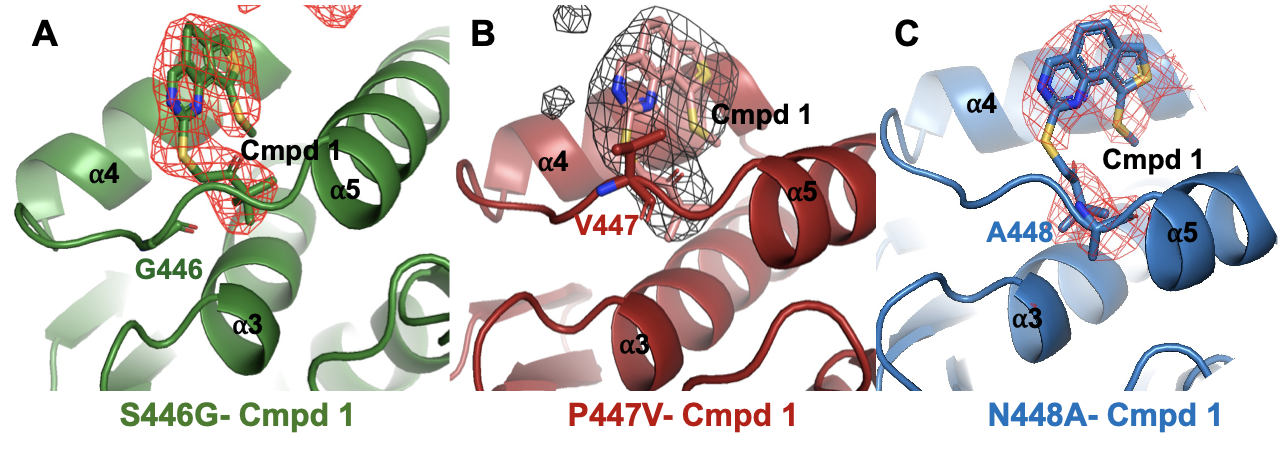

**Figure S11**. **Electron density of Cmpd 1 binding to S446G, P447V and N448A variants.** Close-up views of Cmpd 1-bound to the allosteric site in *A*, S446G (green) *B*, P447V (dark red) and *C*, N448A (blue) highlighting the position of the compound relative to the α3–α5 helices. The omit electron density maps (Fo–Fc, contoured at 3.0σ) are shown as a mesh, validating the presence and positioning of Cmpd 1 in each complex. The figures were generated using the PyMol visualization tool.
